# Signature molecular changes in the skeletal muscle of hindlimb unloaded mice

**DOI:** 10.1101/2020.10.30.362160

**Authors:** M. Azeem, R. Qaisar, A. Karim, A. Ranade, A. Elmoselhi

**Affiliations:** University of Sharjah

## Abstract

Hind-limb unloaded (HU) mouse is a well-recognized model of muscle atrophy; however the molecular changes in the skeletal muscle during unloading are poorly characterized. We have used Fourier transform infrared (FTIR) and Raman spectroscopy to evaluate the structure and behavior of signature molecules involved in regulating muscle structural and functional health. The FTIR and the Raman spectroscopic analysis of gastrocnemius muscles was compared between 16-18 weeks old HU c57Bl/6J mice and ground-based controls. The molecular components of the samples were identified by using the FTIR spectra from the control and the unloaded samples. The Raman spectra showed that the signals for asparagine and glutamine were reduced in HU mice, possibly indicating increased catabolism. The peaks for hydroxyproline and proline were split, pointing towards molecular breakdown and reduced tendon repair. We also report a consistently increased intensity in > 1300 cm^−1^ range in the Raman spectra along with a shift towards higher frequencies in the HU mice, indicating activation of sarcoplasmic reticulum (SR) stress during HU.

**SIGNIFICANCE:** Mouse model of hindlimb unloading recapitulates many features of disuse muscle atrophy due to spaceflight and prolonged bed rest. However, a thorough understanding of molecular changes underlying muscle detriment in disuse partly remains elusive. We have applied the spectroscopic and density functional techniques conjointly to characterize the molecular changes in the skeletal muscle of hindlimb unloaded mice. A number of conformational changes and the breakdown of the molecular bonds in the skeletal muscle are observed, which can potentially dictate loss of muscle mass and strength during mechanical unloading. Our reporting of the signature spectral changes in the unloaded skeletal muscle can be a useful step towards the therapeutic interventions targeting specific molecules.

## INTRODUCTION

Mechanical unloading of the skeletal muscle leads to a rapid loss of the muscle mass and the strength, which worsens with increasing duration of unloading. This condition is relevant to a plethora of scenarios from prolonged bed rest due to stroke, chronic diseases and bone fractures (1) as well as spaceflights by the astronauts (2). However, the search for an effective pharmacological therapy as a countermeasure remains elusive, partly because the molecular mechanisms of muscle detriment in unloading remain poorly understood. A detailed molecular mapping of the skeletal muscle in unloading conditions is required to design therapeutic drugs with specific molecular targets. Over the past three decades, a large number of studies have investigated the molecular changes during muscle unloading. However, a rigorous characterization of these changes has not been performed. The spectroscopy offers a unique non-biological approach to obtain a snapshot of global molecular changes in skeletal muscle during mechanical unloading.

The techniques of the FTIR and the Raman spectroscopy has effectively been applied to the biological tissues (3–10), including for cancer diagnosis (11, 12). The characteristic peaks in the spectroscopic data of the biological samples are associated with the vibrational frequencies of certain molecules in the sample. However, the interpretations of the spectra in the literature are not consistent, partly because of variations in sample storage and preparation as well as the type of the techniques used.

In particular, the intrinsic flexibility of the biological molecules makes the interpretations of the Raman spectra more challenging. For example, the protein molecules in the biological tissues are dynamic, adopting different spatial structures and geometries in various conditions. Each geometrical configuration of a molecule has its own unique vibrational fingerprint, and the actual spectrum is the total of the fingerprints of all spatial molecular variants of a molecule. In addition, a certain low-energy molecular configuration is more stable than a high-energy one, which implies that not all the variants of a molecule are equally represented in a spectrum. This structural variation poses a significant problem for the characterization of the molecules through spectroscopic techniques.

In order to minimize these errors, we have adopted a unique approach of conjoining the theoretical and experimental techniques. The FTIR was used to identify the molecules present in the sample. The theoretical Raman spectra for the molecules was, then, calculated by performing the quantum mechanical density functional theory (DFT) calculations. The quantum mechanical approach can also be used to accurately describe the polarization and the charge transfer effects in the biologically active molecules due to subtle chemical reactions. By using the experimental and theoretical techniques, we have aimed to characterize the preferential molecular changes in the skeletal muscle under unloading conditions using the HU mouse model. In evaluating the molecular phenotypes of biological tissues, our findings unravel novel molecular changes in the skeletal muscle under unloading conditions.

## MATERIALS AND METHODS

Male, c57BL/6J mice were maintained under pathogen-free conditions, housed 2-3 mice per cage (control) or one mouse per cage (unloaded) in a 12:12 (light: dark) cycle and provided with food and water *ad libitum,* as described elsewhere (13). All mice were 16-18 weeks old at the start of the experiments. One group of mice was mechanically unloaded with hind limbs suspended in air, while the other group was kept as the ground-based controls for a similar duration for 15 days. After the period of unloading, mice were euthanized via cervical dislocation. Gastrocnemius muscle tissues were collected from the control and unloaded mice (N = 4-5 / group) and were immediately snap-frozen in liquid nitrogen for analysis via FTIR and Raman Spectroscopy as shown schematically in Fig. 1.

**Fig. 1.**
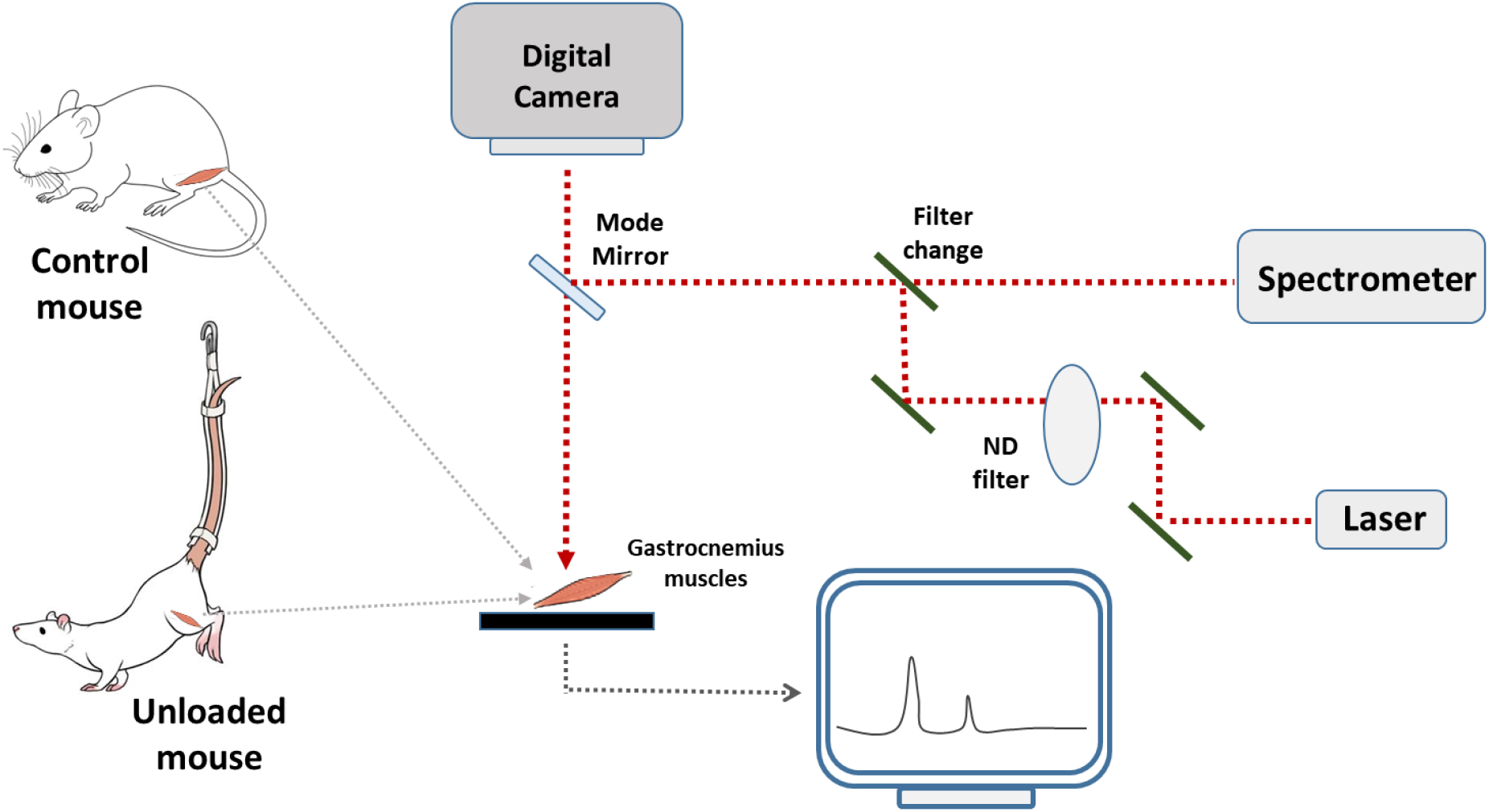
Schematic representation of the experimental procedures.

The FTIR spectra was collected by using JASCO FTIR model 6300. The number of scans for a sample was 20 and the scan time was 35 seconds. Resolution for the FTIR spectra was 2.0 cm^−1^ and the spectral range was from 800 to 2000 cm^−1^. The Raman spectra were obtained by using Renishaw in via Raman spectrometer. Three specimens were selected for the control and the unloaded samples each and 10 spectra were collected from various locations of each muscle to obtain the average. In each recording, a site of a specimen was exposed to a 785 nm wavelength laser with a spot size of 50 μm and a laser power of 1%. The spectral range was from 100 cm^−1^ to 1700 cm^−1^, the fingerprint region where the signature peaks from the biological molecules can be found. To avoid damage to the muscle tissues from the intensity of the laser, the spectra were collected at the intervals of 500 cm^−1^ and then stitched together to obtain the full spectrum. The acquisition time for individual spectra was 10 seconds.

## THEORY AND CALCULATIONS

The frequencies and the intensities of the spectra were fitted with the Lorentzian function (14) to resolve peaks contributed by each molecule in the sample. The Lorentzian peak function is define as,

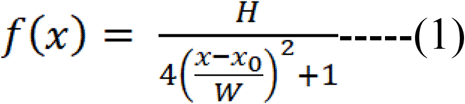

where *x*_*0*_, *H* and *W* are the peak position, the height of the peak and the full width half maximum (FWHM), respectively.

The natural shape of a spectral line is Lorentzian, which has a sharp narrow peak with most of the intensity of the peak located in the tails extending to infinity. The uncertainty in the excitation response, the location of the energy level of the excited state of the molecule as well as the duration for which it remains in the excited state influence the width of the Lorentzian line. Due to the scattered nature of the experimental spectra, the uncertainty in the position of peaks produced by the Lorentz fit is up to ± 5 cm^−1^. All the spectra were normalized with respect to the strongest peak.

For the theoretical calculation of the Raman spectra, the choice of exchange and correlation functionals is important in DFT calculations. In this work, B3LYP functional was used, which combines Slater, Hartree–Fock and Becke (15) exchange with the correlation terms due to Lee, Parr and Yang (16). It is a global hybrid generalized gradient approximation functional (GH-GGA), which is applied to determine the electronic structure of the biological systems (17). A 6-31G^*^ polarization basis set was chosen for the density functional models. A molecular vibration is Raman active, if it changes the polarizability of the molecule. Therefore, a B3LYP/6-31-G* is the most suitable approximation to calculate the Raman spectra for the biological systems. All calculations were performed by using Spartan’18 Parallel Suite, Version 1.3.0 (18).

The data about muscle strength, mass and mRNA concentration is presented using mean and standard error of mean. Student’s *t*-test was used for comparison between the two groups. A p-value < 0.05 was considered to be statistically significant. Statistical analysis was performed using GraphPad Prism version 8.

## RESULTS

The FTIR spectra arise due to the absorption of infrared light and mainly affects the dipole bands such as carbon-oxygen double bods, single bonds of oxygen-hydrogen and nitrogen-hydrogen (19). The FTIR spectra collected from the control and unloaded samples are shown in the Fig. 2. The band positions for both groups of samples are almost the same. The band at 1042 cm^−1^ is the result of the coupled stretching and bending vibrations of the C-H and C-O and C-OH groups of carbohydrates, sugar and starch. Similar vibrational modes are present at 1062 cm^−1^; however these are contributed by the tyrosine molecules of proteins. The bending vibrations of the C-H and O-H groups of tryptophan molecules appear at 1207 cm^−1^. Amide III bands are visible in the region of 1200 cm^−1^ to 1350 cm^−1^, which involves vibrations due to C=O, C-N and N-H amide bonds. These are the characteristic features of the proteins (20). The band at 1400 cm^−1^ is the contribution of the symmetric CH3 bending of the methyl groups in proteins. The C-H ring stretching of the tyrosine and tryptophan molecules, as well as N-H wagging vibrations are also contributors to this band. Symmetric CH2 vibrations of tyrosine, hydroxyproline and proline give rise to the bands at 1456 cm^−1^. The well-known amide II bands are present between 1500 cm^−1^ and 1600 cm^−1^. In this region, the stretching of the strong C=N bonds along with N-H bending in the asparagine molecule gives rise to bands at 1635 cm^−1^ and 1642 cm^−1^. A weak coupling of the same modes of vibrations in the glutamine also contributes at 1635 cm^−1^. Finally, weaker amide I bands are present at 1720 cm^−1^ and 1742 cm^−1^. These bands are the results of a weak coupling C=O and C-N stretching and N-H bending. It is interesting to note that the amide I region is more pronounced in the control samples.

**Figure 2.**
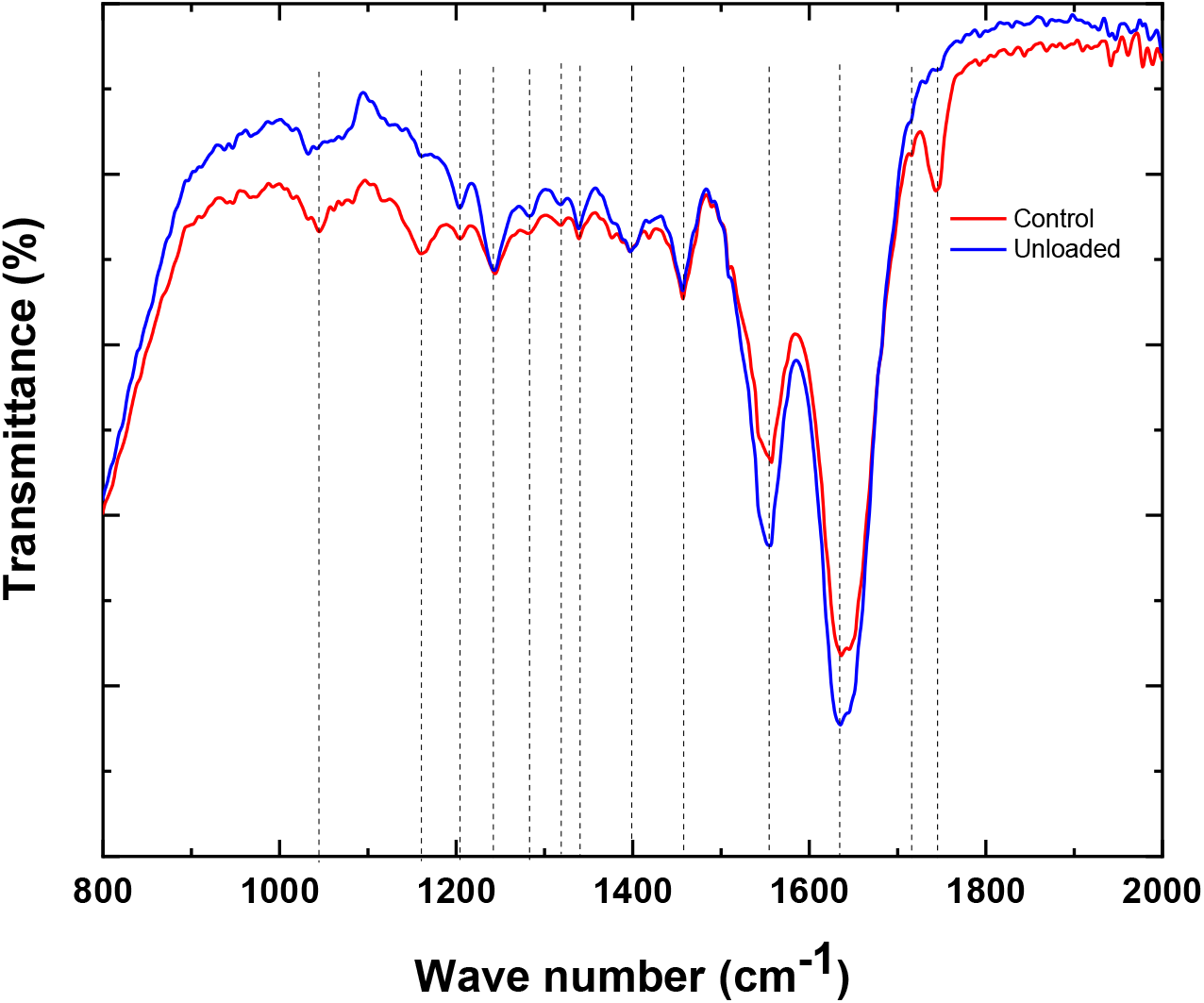
The FTIR spectra from the control and unloaded samples.

The Raman spectra, on the other hand, are the effect of the emission of the scattered light due to the change in the polarizability of the covalent bonds. Therefore, any signature changes registered in the control and unloaded samples can be read by the Raman spectroscopy. The Raman spectrum from a control sample is shown in Fig. 3 (A) along with a Lorentz fit. The scattered spectrum indicates that the peaks contributed by the different molecules overlap. The Lorentz fit, therefore, resolved the peaks more clearly to help identify the types and structures of the molecules as well as their vibrational modes.

**Figure 3.**
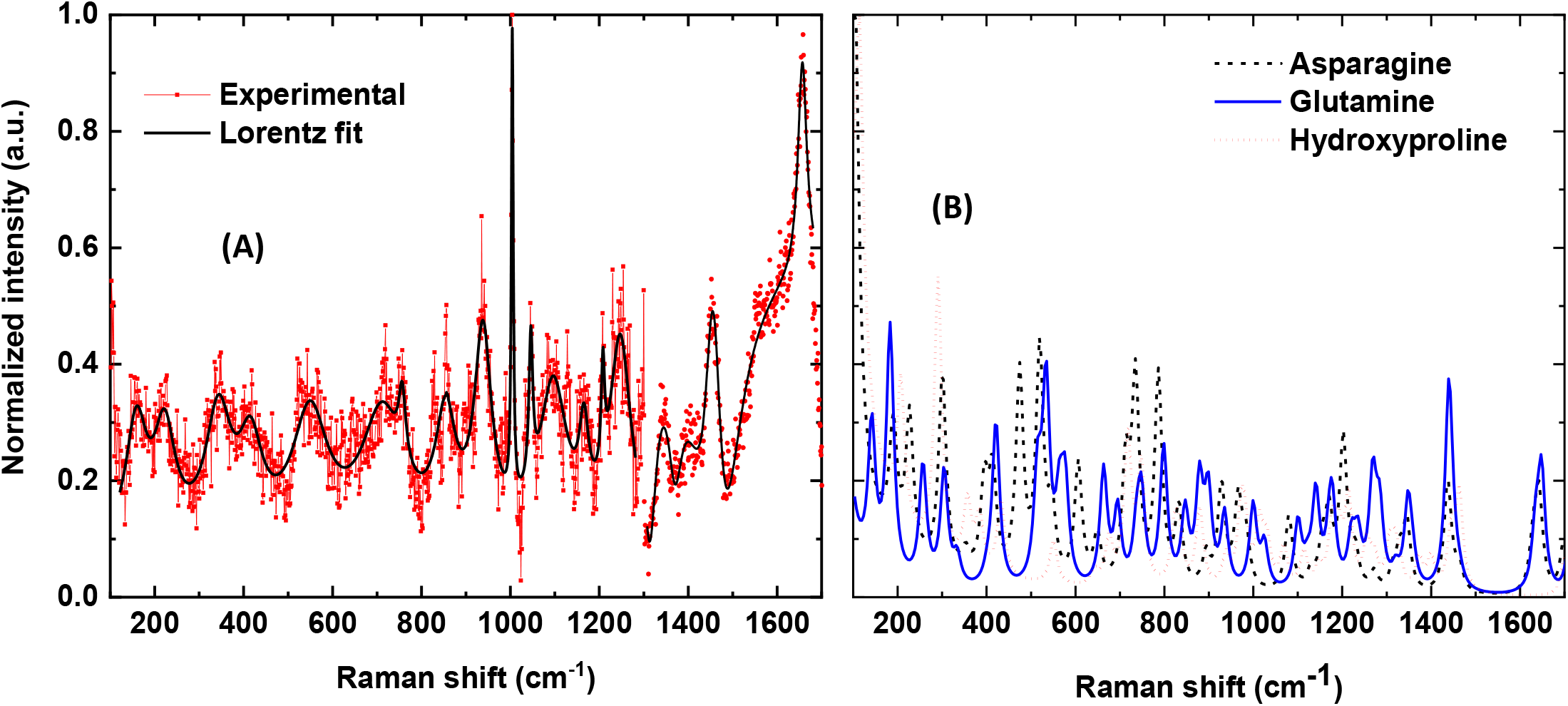
(A). The experimentally obtained Raman spectrum of control specimen along with Lorentz fit. (B) The calculated Raman spectra for the Asparagine, glutamine and the hydroxyproline molecules.

The theoretical Raman spectra were calculated for asparagine, glutamine, hydroxyproline, DL-tyrosine, UDP-D-glucose, praline, proline and tryptophan as these are the most likely molecules present in the specimen identified by the FTIR spectra (Fig. 2). For the sake of clarity, only the calculated spectra from the asparagine, glutamine and hydroxyproline are shown in Fig. 3B. Our experimental and the calculated Raman peaks agree well with the previous studies (6, 21, 22) within the uncertainty. However, the calculated spectra also show additional Raman bands which are, either very weak or indistinguishable in the experimentally obtained spectra. It should be noted that the calculations were carried out in the gas phase on single molecules with an ideal spatial geometry, which may have caused the extra Raman bands to appear in the calculated spectra. In the actual samples, the quantum states of different molecules are entangled suppressing certain vibrational modes while enhancing the others. Table 1 lists the calculated band positions for all the molecules.

**Table 1.** The calculated Raman band positions for molecules in the skeletal muscles.

| Molecule | Calculated Band positions (cm<sup>-1</sup>) |
| --- | --- |
| Asparagine | 190, 227, 301, 353, 395, 414, 474, 353, 395, 414, 474, 519, 547, 606, 649, 713, 735, , 787, 832, 894, 928, 966, 983, 1080, 1130, 1156, 1170, 1202, 1229, 127, 1314, ,1329, 1345, 1437, 1463, 1630, 1642, 1698 |
| Glutamine | 142, 183, 257, 305, 333, 402, 421, 515, 534, 564, 577, 663, 695, 737, 748, 798, 847, 879, 899, 935, 1000, 1025, 1100, 1122, 1141, 1170, 1177, 1221, 1235, 1269, 1284, 1320, 1348, 1358, 1440, 1449, 1464, 1635, 1648 |
| Hydroxyproline | 114, 207, 291, 357, 384, 432, 434, 550, 644, 676, 718, 748, 823, 833, 879, 939, 973, 981, 1014, 1031, 1069, 1114, 1148, 1177, 1192, 1214, 1255, 1272, 1306, 1318, 1329, 1373, 1400, 1445, 1463, 1700 |
| DL Tyrosine | 160, 193, 275, 282, 300, 331, 378, 392, 405, 413, 457, 492, 515, 573, 628, 637, 689, 727, 748, 763, 798, 821, 824, 847, 884, 912, 920, 979, 987, 1068, 1081, 1135, 1145, 1147, 1169, 1182, 1208, 1255, 1268, 1280, 1300, 1316, 1322, 1373, 1416, 1444, 1494, 1587, 1598, 1603 |
| UDP-D-Glucose | 126, 132, 162, 171, 227, 235249, 254, 282, 303, 376, 424, 520, 554, 578, 587, 608, 710, 714, 750, 753, 792, 1207, 1218, 1230, 1239, 1253, 1255, 1275, 1289, 1454, 1612, 1667 |
| Praline | 174, 204, 252, 285, 298, 464, 484, 487, 492, 529, 637, 710, 799, 837, 966, 1023, 1028, 1142, 1194, 1214, 1321, 1369, 1383, 1437, 1444, 1612, 1680, 1688 |
| Proline | 168, 215, 314, 387, 474, 526, 620, 699, 775, 825, 880, 920, 942, 980, 1063, |
|  | 1078, 1090, 1115, 1181, 1207, 1227, 1269, 1290, 1301, 1314, 1330, 1408, 1451, 1466, 1480 |
| Tryptophan | 160, 551, 736, 745, 750, 1006, 1331, 1349, 1420, 1450, 1518, 1603 |

By comparing our results with the previous studies(6, 21, 22) and the calculated Raman spectra, the bands at 159 cm^−1^, 223 cm^−1^, 344 cm^−1^, and 1209 cm^−1^ were allocated to asparagine, whereas the bands from asparagine and glutamine overlapped at the frequencies 412 cm^−1^, 533 cm^−1^ and 551 cm^−1^. The peaks at 710 cm^−1^ and 755 cm^−1^ were due to the combined contributions of the hydroxyproline, UDP-D-glucose and tryptophan. A peak at 1004 cm^−1^ is also assigned to the tryptophan molecule. A contribution from the tyrosine molecule was at 595 cm^−1^ and the proline signature bands were at 936 cm^−1^, 1046 cm^−1^, 1090 cm^−1^, 1125 cm^−1^, 1166 cm^−1^, and 1245 cm^−1^. The bands at 1343 cm^−1^ and 1450 cm^−1^ were due to the twisting and bending of the CH3 and CH2 bonds of praline and proline molecules. Similarly, shoulders at 1557 cm^−1^ and 1645 cm^−1^ were the signatures of carbon-carbon bond stretching in the praline molecule.

Next, we compared the Raman spectra of the control and unloaded samples as shown in Fig. 4. We found major differences in the positions and intensities of both the spectra, indicating qualitative and quantitative changes in the molecular cohorts constituting skeletal muscles. The peak at 159 cm^−1^ for the control sample is shifted to the lower frequency of at around 151 cm^−1^ in the unloaded sample, while the intensity of the peak at 220 cm^−1^ is slightly increased in the unloaded sample. The intensities of the unloaded sample at the 344 cm^−1^ and 412 cm^−1^ are significantly reduced compared to the control along with the appearance of a new peak at 460 cm^−1^, indicating conformational changes in the asparagine and the glutamine molecules. These amino acids are involved in regulating a variety of functions in the skeletal muscle including bioenergetics and protein synthesis (23). Consequently, changes in their molecular orientation and/or relative proportions in the HU skeletal muscle can be potential contributors to the muscle atrophy and weakness in these mice.

**Figure 4.**
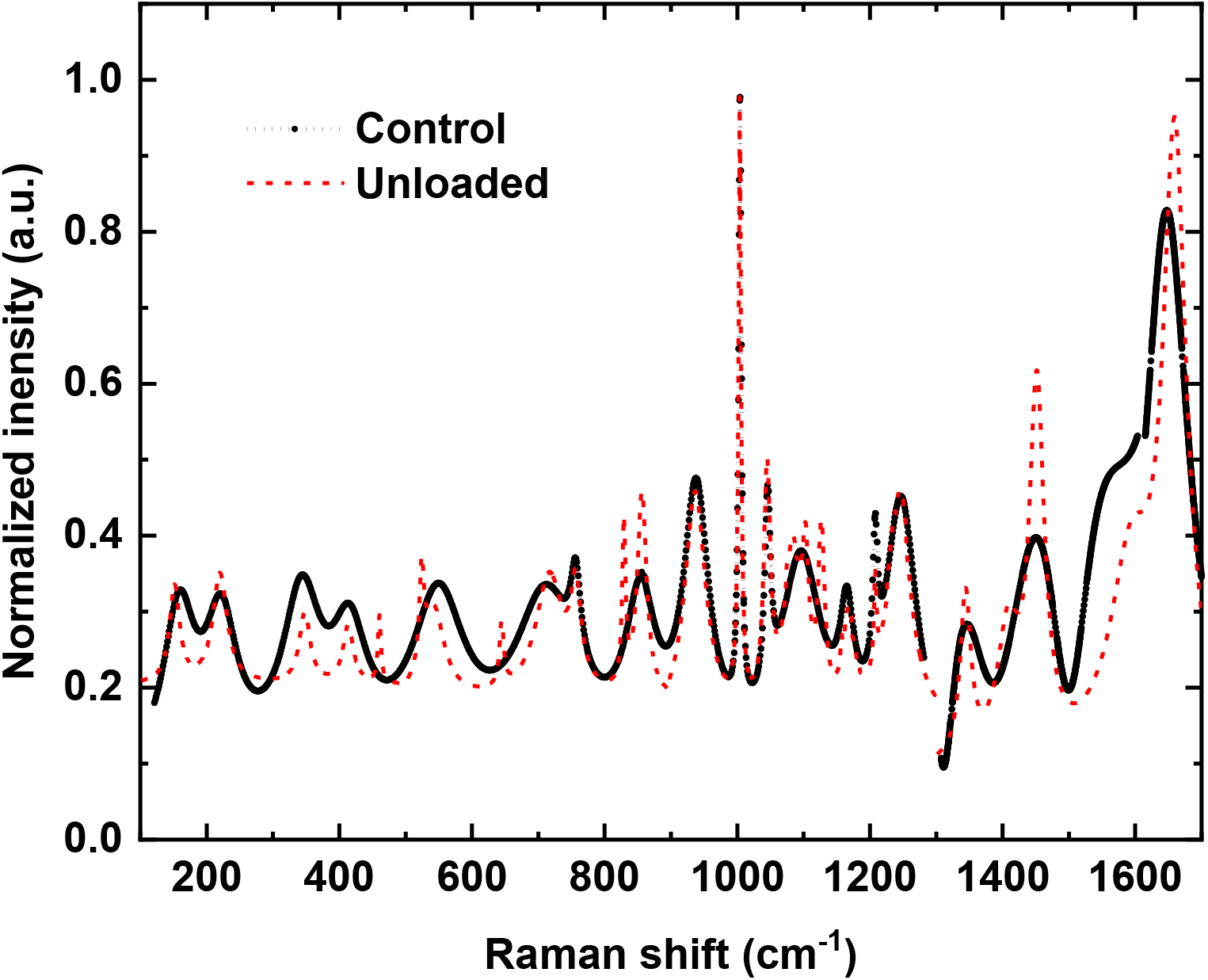
Experimentally obtained Raman spectra from a control and unloaded specimen.

We also found that the peak at 551 cm^−1^ was slightly shifted to the lower frequency of 525 cm^−1^, with a shoulder appearing at 542 cm^−1^ for the asparagine and glutamine molecules. Additionally, we found a weak but sharp peak at 644 cm^−1^ in the unloaded samples only, which was most probably due to hydroxyproline. The hydroxyproline is a derivative of proline and is a structural constituent of multiple muscle proteins. It is found in collagens in high concentrations and has a supportive role during muscle contraction. Other hydroxyproline peak at 709 cm^−1^ was slightly shifted to a higher frequency of 717 cm^−1^ in the unloaded sample along with a small decrease in the intensity of the peak at 756 cm^−1^. The changes in the height and displacement of the signature peak of hydroxyproline potentially means altered quality and/or quantity of this amino acid, which can have pathological consequences in the conditions of prolonged HU. The peak associated with tyrosine at 856 cm^−1^ was also split into two sharp and narrow peaks at 829 cm^−1^ and 857 cm^−1^ with higher intensity in the HU mice, when compared to control mice. Tyrosine is an alpha amino acid and plays an important role in the enzymatic activities of many proteins. Further, the proline peak at 1090 cm^−1^ was also split into three narrow peaks (triplet) at 1087 cm^−1^, 1103 cm^−1^ and 1128 cm^−1^. A doublet in the tyrosine and a triplet in the proline molecules suggests that some molecular residues in proteins undergo a change in their hydrogen bonding environment (24). Thus, the changes in the tyrosine and proline can potentially contribute to the loss of muscle mass and/or strength in the unloading conditions.

The spectra above 1300 cm^−1^ showed some interesting findings. The spectral intensities of the unloaded muscles in this range were much stronger than the control muscles at all the positions with a shoulder forming at 1410 cm^−1^. The intensity of the shoulder at 1557 cm^−1^ was significantly reduced and shifted to 1599 cm^−1^ for the unloaded samples. Similarly, the peak at 1645 cm^−1^ was also shifted to 1659 cm^−1^.

## DISCUSSION

It is apparent that the unloaded samples have catabolic changes at the molecular level, with biological implications. The reduction in the Raman signal intensities from asparagine and glutamine indicate an enhancement of the catabolic processes in unloaded muscle. The splitting of the hydroxyproline and proline peaks signifies the breakdown of these molecules, which is most likely associated with reduced healing and repair processes in the skeletal muscle during mechanical unloading. Similarly, a breakdown in the tyrosine molecules indicates reduced protein synthesis, which can potentially contribute to muscle atrophy and weakness. In accordance with the changes in Raman spectral analysis, we found significant reduction of the gastrocnemius muscle mass and grip force in HU mice, when compared with the control, in accordance with our previous findings (Fig. 5A) (13). This can partly be due to the changes taking place in the frequency range of 100 cm^−1^ to 1300 cm^−1^, according to the Raman spectra.

**Fig. 5.**
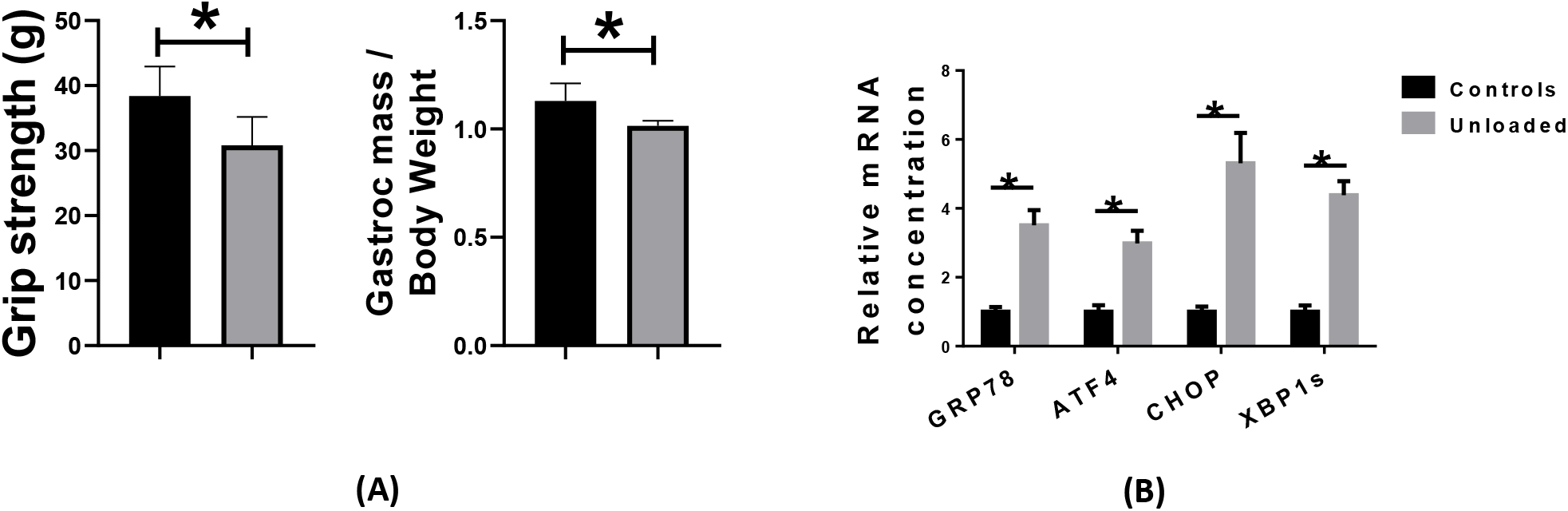
(A) The gastrocnemius muscle mass and grip strength in the control and unloaded mice, *p < 0.05. (B) Relative mRNA concentrations of the markers of ER stress in gastrocnemius muscles of control and unloaded mice. (N = 5-8 per group), *p < 0.05.

The changes in the spectral range greater than 1300 cm^−1^ have been attributed to the accumulation of unfolded protein in the sarcoplasmic reticulum (SR), a condition called SR stress (25). In order to perform an independent validation of these findings, we conducted RT-PCR experiments as described previously (26). Specifically, we quantified the signature mRNA markers of SR stress in the gastrocnemius muscles of control and unloaded mice. A significant upregulation in the mRNA expressions of GRP78, ATF4, CHOP and s-XBP-1 was reported in the unloaded mice (Fig 5B,), which indicate heightened SR stress and validates the Raman spectroscopic findings.

The SR stress is emerging as an important candidate process to the muscle decline during various catabolic conditions (27) and can potentially contribute to the loss of muscle mass and strength reported here in the HU mice. Thus, the perturbation of SR protein folding capacity, as indicated by changes in the spectral range > 1300cm^−1^, seems a prime contributor to muscle detriment in the HU mice. The enhanced intensity of the Raman spectra in this range in the unloaded samples is also accompanied by a shift towards the higher frequency, a result of the broken molecules with shorter bond lengths, thus shifting the Raman peaks to the higher wave numbers. In addition, as described above, the spectra in the range of 1300 cm^−1^ to 1700 cm^−1^ are mainly the result of the CH2, CH3 vibrations; therefore the higher intensity of the unloaded samples in this energy range was due to the accumulation of the broken molecules.

## CONCLUSION

We have investigated the signature molecular changes in the skeletal muscles of the HU mice by using the Raman spectrometer. We conclusively show that the muscle atrophy and weakness in the unloaded samples are associated with the significant changes in the molecular phenotypes of skeletal muscle. For the first time, the experimental and the theoretical Raman spectra are conjoined to study the molecular constituents and to track the changes in them. Our key findings are the significant differences in the features of the Raman spectra between control and unloaded mouse muscles. The major components of the skeletal muscles are asparagine, glutamine, hydroxyproline, DL tyrosine, UDP-D-glucose, proline, praline and tryptophan. The concentrations of the asparagine and the glutamine molecules were significantly changed in the unloaded muscles as evident by the reduction in the intensities of the Raman peaks at 344 cm^−1^ and 412 cm^−1^. The peaks at 460 cm^−1^, 525 cm^−1^, 542 cm^−1^ and 551 cm^−1^ in the unloaded samples illustrate the conformational changes in these molecules. The signatures peaks of hydroxyproline were at 709 cm^−1^ and 756 cm^−1^ in the control muscles, while for the unloaded muscles, a new peak appears at 644 cm^−1^ indicating the breakdown of the molecular bonds in the hydroxyproline. Similarly, the splitting and the shifting was observed in the peaks associated with tyrosine and the proline molecules. The spectra from the control and unloaded samples was also clearly distinguishable in the spectral ranges greater than 1300 cm^−1^ indicative of heightened SR stress, which was independently validated by RT-PCR data.

## AUTHORS CONTRIBUTION

M. A. collected the experimental data from Raman and FTIR spectrometer, applied DFT theory and wrote the manuscript. R. Q. prepared the samples, performed statistical analysis and collected RT-PCR data and wrote the parts of the manuscript. A. R., A. K., and A. M. provided assistance to R. Q. in the sample preparation and storage.

## ACKNOWLEDGEMENT

The work is supported by the UOS competitive research grant 1802143076. Authors are thankful to Dr. Hussain Alawadhi of centre of advanced materials research, UOS, to provide unlimited access to the spectrometers.

